# unCOVERApp: an interactive graphical application for clinical assessment of sequence coverage at the base-pair level

**DOI:** 10.1101/2020.02.10.939769

**Authors:** Emanuela Iovino, Marco Seri, Tommaso Pippucci

## Abstract

**Motivation:** Next Generation Sequencing (NGS) is increasingly adopted in the clinical practice largely thanks to concurrent advancements in bioinformatic tools for variant detection and annotation. Despite improvements in available approaches, the need to assess sequencing quality down to the base-pair level still poses challenges for diagnostic accuracy. One of the most popular quality parameters of diagnostic NGS is the percentage of targeted bases characterized by low depth of coverage (DoC). These regions potentially hide a clinically-relevant variant, but no annotation is usually returned for them.

However, visualizing low-DoC data with their potential functional and clinical consequences may be useful to prioritize inspection of specific regions before re-sequencing all coverage gaps or making assertions about completeness of the diagnostic test.

To meet this need we have developed unCOVERApp, an interactive application for graphical inspection and clinical annotation of low-DoC genomic regions containing genes.

**Results:** unCOVERApp is a suite of graphical and statistical tools to support clinical assessment of low-DoC regions. Its interactive plots allow to display gene sequence coverage down to the base-pair level, and functional and clinical annotations of sites below a user-defined DoC threshold can be downloaded in a user-friendly spreadsheet format. Moreover, unCOVERApp provides a simple statistical framework to evaluate if DoC is sufficient for the detection of somatic variants, where the usual 20x DoC threshold used for germline variants is not adequate. A *maximum credible allele frequency* calculator is also available allowing users to set allele frequency cut-offs based on assumptions about the genetic architecture of the disease instead of applying a general one (e.g. 5%). In conclusion, unCOVERApp is an original tool designed to identify sites of potential clinical interest that may be hidden in diagnostic sequencing data.

**Availability:** unCOVERApp is a freely available application written in R and developed with Shiny packages and available in GitHub.

## 1 Introduction

In the span of less than a decade, Next Generation Sequencing (NGS) has established itself as the gold-standard approach for the molecular diagnosis of genetic diseases (Biesecker et al, 2014). In the diagnostic setting, NGS allows for the simultaneous inspection of all the genes underlying a genetically heterogeneous condition, considerably reducing analysis costs and times to diagnosis.

One of the most popular quality parameters of diagnostic NGS is the percentage of targeted bases with depth of coverage (DoC) above a threshold considered adequate for accurate variant identification (usually 20x). However, sequence DoC may drop below this threshold due to technical issues in hybrid capture or alignment even across clinically-relevant genes (Sanghvi et al, 2018). Diagnostic laboratories may use visual inspection of each low DoC region and Sanger sequencing to fill in the DoC gaps. Especially when the gene set is large (including up to hundreds of genes) this task may become tedious and time-consuming. Alternatively, limitations of the test due to low DoC regions should be clearly stated in the diagnostic report (Matthjis et al 2016).

A vast range of functional predictions and clinical annotations are made available in genomic databases such as dbNSFP (Liu X et al., 2016). NGS bioinformatic pipelines draw information from these databases to contextualize variants within a functional and clinical frame.

Conversely, no annotation is returned for non-variant sites. If affected by low DoC, these positions potentially harbor a “hidden” variant. Tools are available (Sanghvi et al, 2018) that select and report regions below a certain coverage threshold in a format easy to export on genome browsers. However, it would be helpful to directly visualize low DoC data and to know potential sequence changes in positions that have an annotated clinical consequence or a predicted deleterious effect. This knowledge may be useful to prioritize inspection of specific regions containing clinically-relevant low DoC positions, instead of immediately filling all the gaps and before making diagnostic assertions.

Here we present unCOVERApp, an application for both visualization and annotation of coverage gaps in NGS data.

## 2 Methods

unCOVERApp is a freely available application under MIT license, written in R language and implemented in Shiny (Chang et al., 2016), a framework that enables to build web applications using R. unCOVERApp provides interactive graphical DoC analysis from whole gene to base-pair resolution using R package Gviz for live genomic annotation (Hahne et al., 2016). To associate low-DoC sites with functional and clinical annotations, unCOVERApp uses dbNSFP version 4.0. Calculator of maximum credible population Allele Frequency (AF) (Whiffin et al., 2016) is integrated in unCOVERApp to allow user-defined AF thresholds. To calculate the adequate DoC for somatic variants, unCOVERApp computes the 95 % probability of the binomial distribution based on an expected allele fraction (probability of success) and a minimum number of variant reads (number of successes).

unCOVERApp takes as input a BED file containing the base-pair DoC across the target space for one or more samples. A companion bash script retrieves DoC from sample BAM file(s) with alignments on either hg19 or hg38 across a user-defined set of genes to prepare the input BED file. unCOVERApp documentation, preparation script and specifications of the required dependencies to run it on a local machine are provided at https://github.com/Manuelaio/unCOVERApp.xs

## 3 Results

unCOVERApp displays three web pages each providing a different analysis module, following the workflow illustrated in Figure 1. An example of unCOVERApp analysis is shown as Supplementary Material.

**Figure 1.**
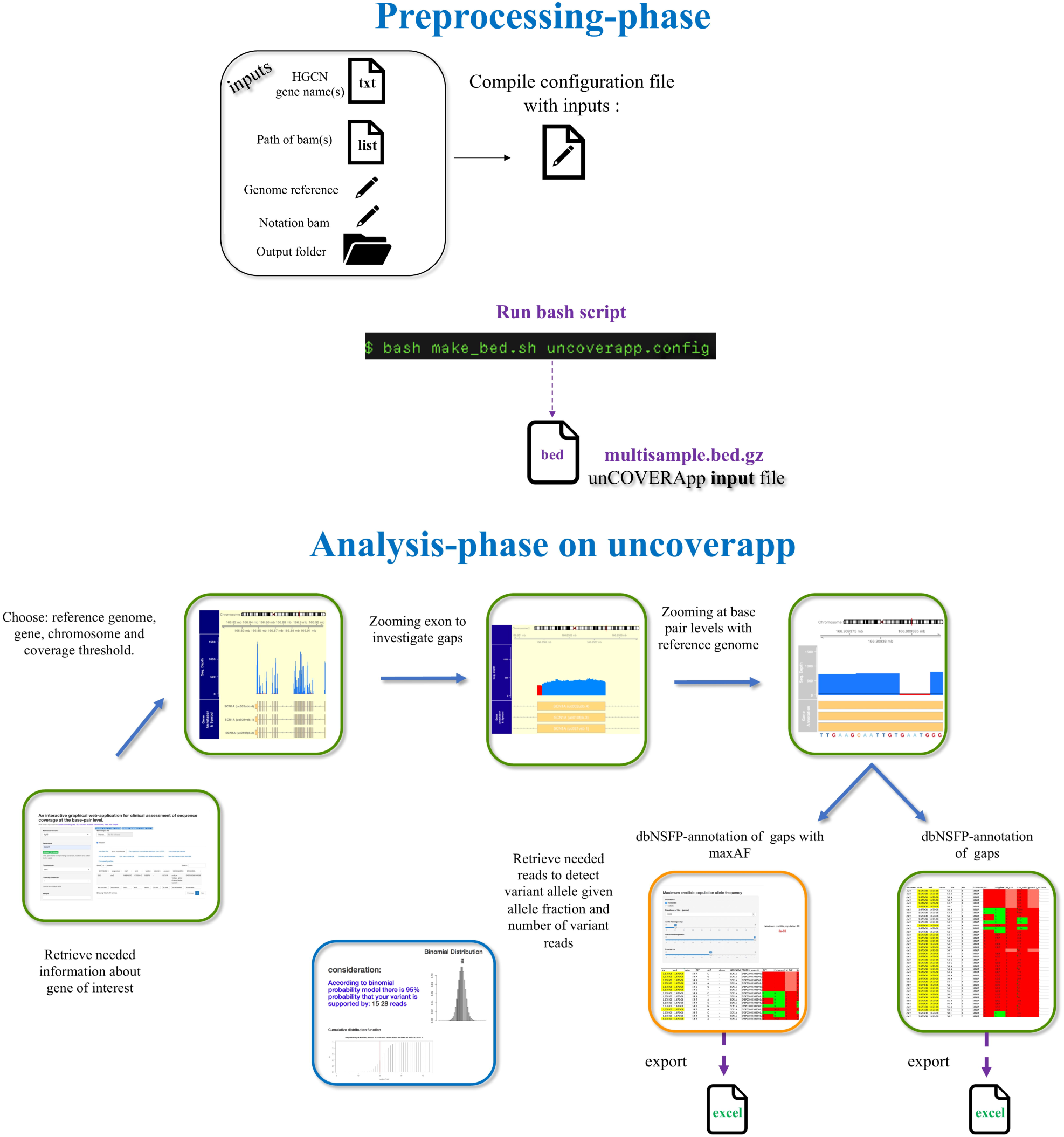
unCOVERApp workflow for sequencing data. The application serves a bash script and a configuration file, as static files, to simply prepare input loading file from BAM file(s) and list of HGNC gene name(s) on local machine. Preprocessing scripts generate a BED-zipped file containing depth of coverage for each genomic position of target genes of inspected samples. Only count reads with base quality and with mapping quality greater than INT are considered. Once uploaded on unCOVERApp, BED file could be visualized and annotated.

The main page requires the user to set the desired DoC threshold, reference genome, gene, transcript of interest and sample for the analysis, in order to generate a DoC histogram where target bases below the threshold are colored in red (Supplementary Figure 3). The histogram plot can be zoomed in to user-selected intervals, from exon to base-pair level.

unCOVERApp provides a table with dbNSFP-based annotation of all potential nucleotide changes across low DoC genomic coordinates, downloadable in Excel format. In the table, changes annotated as clinical, high impact or deleterious are highlighted to allow instant evaluation of annotations associated with variants that have been potentially missed due to low-DoC. Example of a pathogenic variant affected by low DoC that was annotated in unCOVERapp is shown in Supplementary Figure 5.

The default AF threshold for a variant to be annotated in unCOVERApp is 5%, recommended as the upper bound for alleles of clinical interest according to the American College of Medical Genetics guidelines (Richards S, 2015). However, a 5% AF threshold is excessive for most disease-related variants. Calculator of the maximum credible population AF (Whiffin N et al., 2016) is available on the second unCOVERApp page. The AF threshold can be modified by the user based on assumptions on the genetic architecture of the disease to automatically adjust the displayed variants.

The general 20x minimum DoC threshold is reasonable for germline events where the expected fraction of variant alleles is around 0.5. Conversely, it is much more questionable for somatic variants as that fraction can be substantially lower. Page 3 of unCOVERApp provides a simple statistical framework to evaluate if DoC is adequate to somatic variant detection at each genomic position where the user can set the allele fraction expected for the disease-related variant and the number of variant reads that the user considers necessary to support variant calling.

## 4 Conclusion

Lack of adequate DoC over sequence positions of clinical interest is likely the most manifest shortcoming that can affect NGS in the diagnostic setting. Most laboratories either perform one or more rounds of Sanger sequencing to fill in gaps or report low DoC regions as a limitation of the test. However, some low DoC positions may have higher chance to harbor a “hidden” pathogenic variant. Obviously, it is not possible to predict the presence of any deleterious variant in low- or no-coverage data (e.g. frame-shift indels). Nonetheless, information about positions that are clinically annotated or are predicted to change into high impact variants (e.g. nonsense substitutions in haploinsufficient genes) can be retrieved. unCOVERApp is a user-friendly visualization and annotation tool, highlighting low DoC positions of potential interest before making any further decision or assertion about the diagnostic completeness of a test.

## Supplementary documentation

### 1 unCOVERApp

unCOVERApp allows users to visualize and annotate low-coverage genomic regions containing genes in sequencing data. In particular, users can obtain:

- interactive graphical DoC analysis from whole gene(s) to base-pair resolution
- clinically and functionally annotations of low-coverage site downloadable in spreadsheet format
- Calculator of maximum credible population Allele Frequency to allow user-defined AF thresholds rather gnomAD gnomAD generic filter.
- 95 % probability of the binomial distribution based on an expected allele fraction (probability of success) and a minimum number of variant reads (number of successes) for somatic variants.

The code of App is available on GitHub here.

### 2 Prerequisites

This app requires following dependencies:

- **samtools v.1.9**
- **R v.3.5.1** or **RStudio**, and run Rscript to set up the environment.

### 3 Instructions

To run locally unCOVERApp, users can clone or download unCOVERApp repository. Annotation files can be downloaded from googledrive and positioned in unCOVERApp directory. The md5sum of the bed and bed.tbi files can be retrieve in repository.

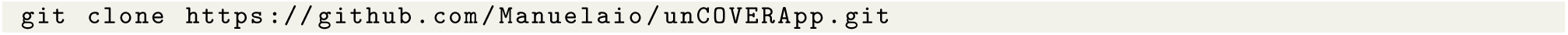

**unCOVERApp** directory must retain the following tree structure.

**Figure 1:**
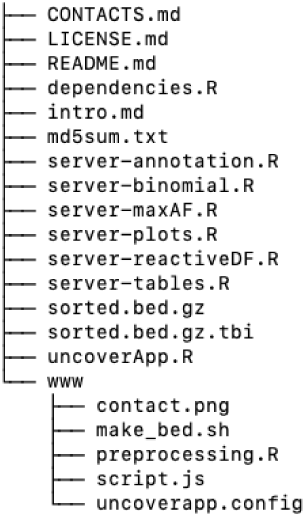
Tree structure of unCOVERApp folder

### 4 Input file preparation

unCOVERApp input file is a BED file (tab-separated) containing depth of coverage (DoC) for each genomic position (one per row) of target genes for as many samples as many BAM files are listed in the “.list” file. In order to easily obtain a input format, users follow the instructions below:

- **write a file with “.txt”** extension containing HGNC gene name(s) (one per row)
- **write a file with “.list”** extension containing absolute path of BAM(s) file (one per row)
- **compile** a configuration file specifying absolute path of: unCOVERApp folder, txt file containing HGNC gene name(s), list file containing absolute path of BAM(s) and folder output location. Compile genome reference and chromosome notation BAM. (number refers to 1, 2, … X,.M notation BAM, chr refers to chr1, chr2,… chrX, chrM notation BAM).

The following image shows an example of compiled configuration file.

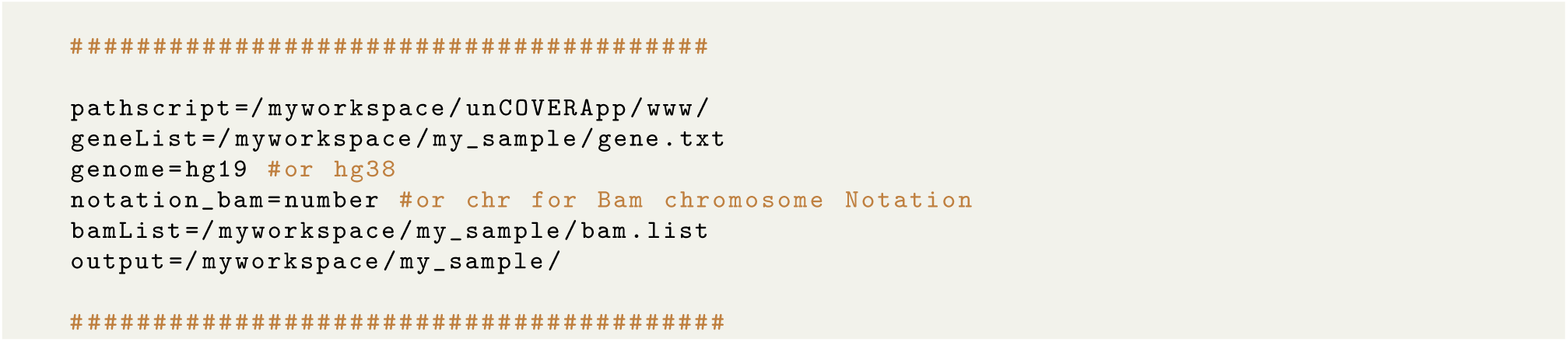

Once users have compiled the configuration file, run the following command through command line

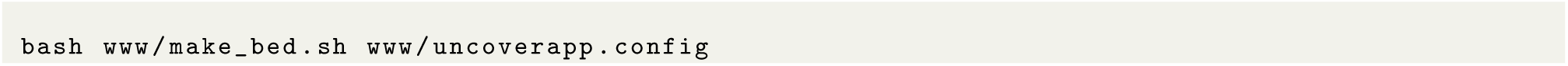

The log file is a trouble shooter, so please revise when any problem happens.

Bash script creates a new directory named with current date in users-defined location, inside is stored input file named **multisample.bed.gz file**.

### 5 unCOVERApp example

Users can run the shiny app with just one command in R:

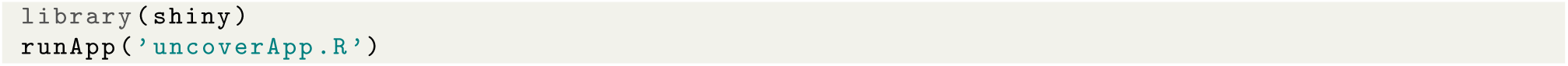

The following example shows how unCOVERApp works and how it helped us to identify pathogenic low coverage position within a candidate gene, POLG, starting from of negative exome sequencing result.

Using bash attached script we have prepared a bed file containing the base-pair DoC across the POLG. We wrote a gene.txt file which contains HGNC official gene name, a file with “.list” extension containing absolute path to BAM sample and we had setup a configuration file.

The first page of unCOVERApp, **Coverage Analysis**, is shown in following figure. Firstly, just made input file was loaded in `Select input box` and visualized in `bed file` table.

Filling inputs as **Reference genome** and **gene name** unCOVERApp returns two outputs:

- `UCSC gene` table, that returns genome coordinates, chromosome number and other useful information based on gene user-defined
- `UCSC exons` table, that returns genome coordinates of each exon for each transcript.

Based on informations provided in outputs, we filled `Chromosome` box and `transcript number` box, moreover we have chosen a coverage threshold and the sample to analyze. Then, unCOVERApp return a plot for graphical inspection of POLG DoC in `Gene coverage` box and a related table with the number of low-coverage positions in each exon given a transcript user-defined. (Figure 3)

**Figure 2:**
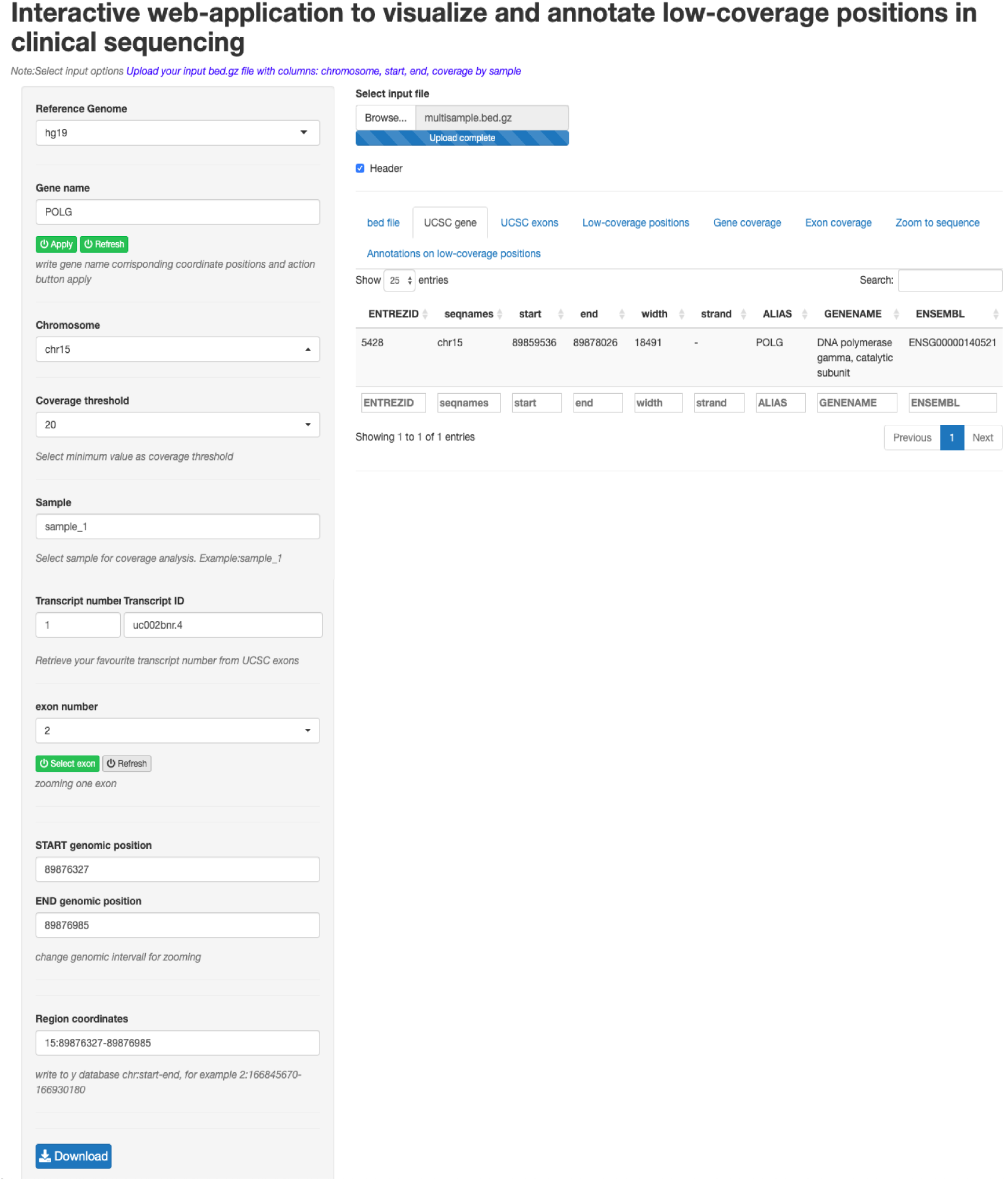
The figure shows first page of unCOVERApp using for coverage analysis in which all required input are filled. As it shows, all required inputs are located in sidebar on the left one by one.

**Figure 3:**
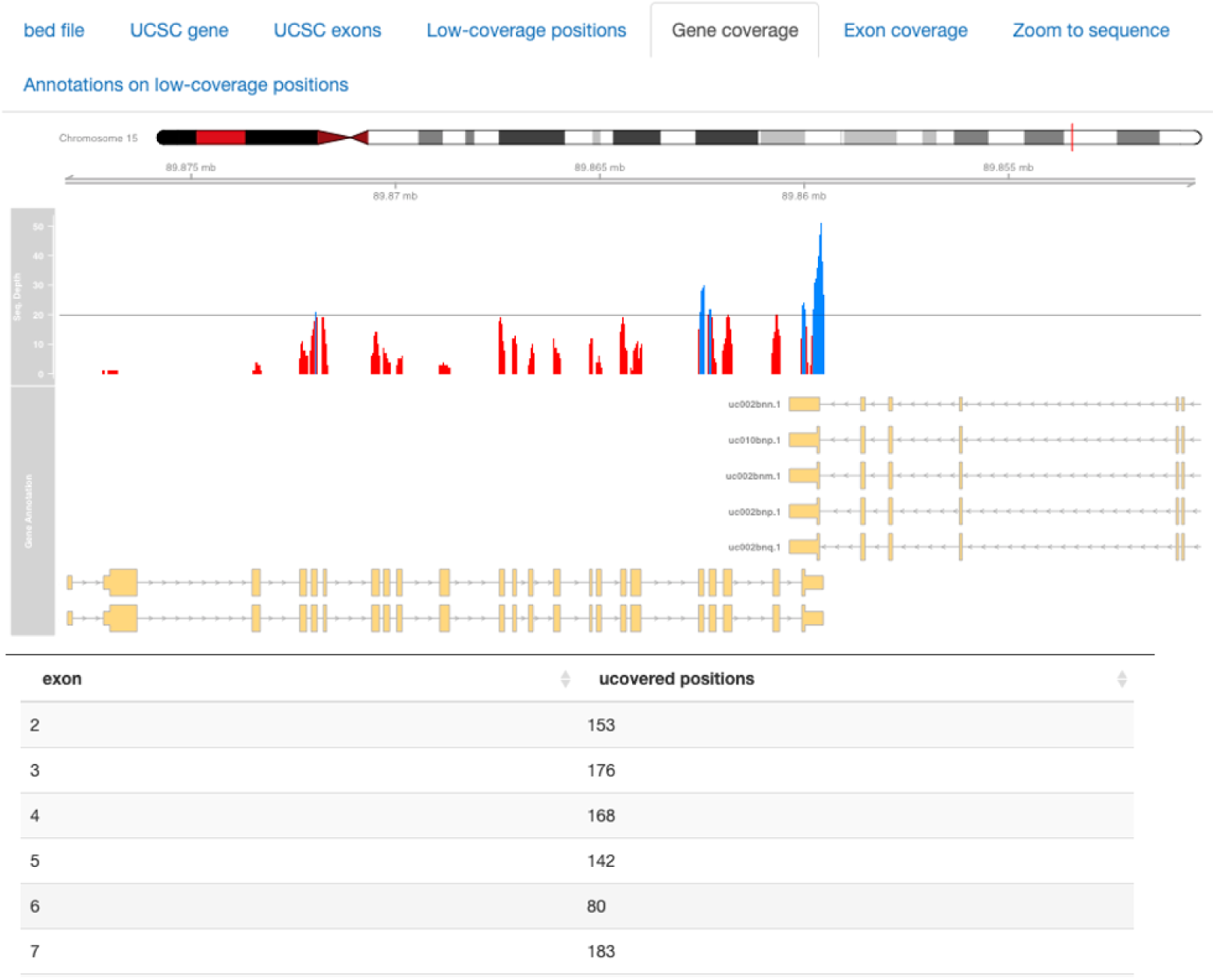
The figure shows DoC of POLG. On top panel it is viewed information about chromosome, genome coordinates and below a DoC information in form of histogram with a dynamically drawn line given a users-threshold cuff off and lastly the different gene transcripts tracks. The table shows the number of uncovered positions for each exon given a chosen transcript.

Table and graphical inspection had shown that the majority of POLG exons are uncovered. Moreover, unCOVERApp provides two zoom function in order to expand the histogram plot in to user-selected intervals, from exon (`exon number` box) to base-pair level (`zoom to sequence`). Inspecting each exon, we have found some low-DoC positions with functional e clinical annotations in exon 10. (Figure 4)

**Figure 4:**
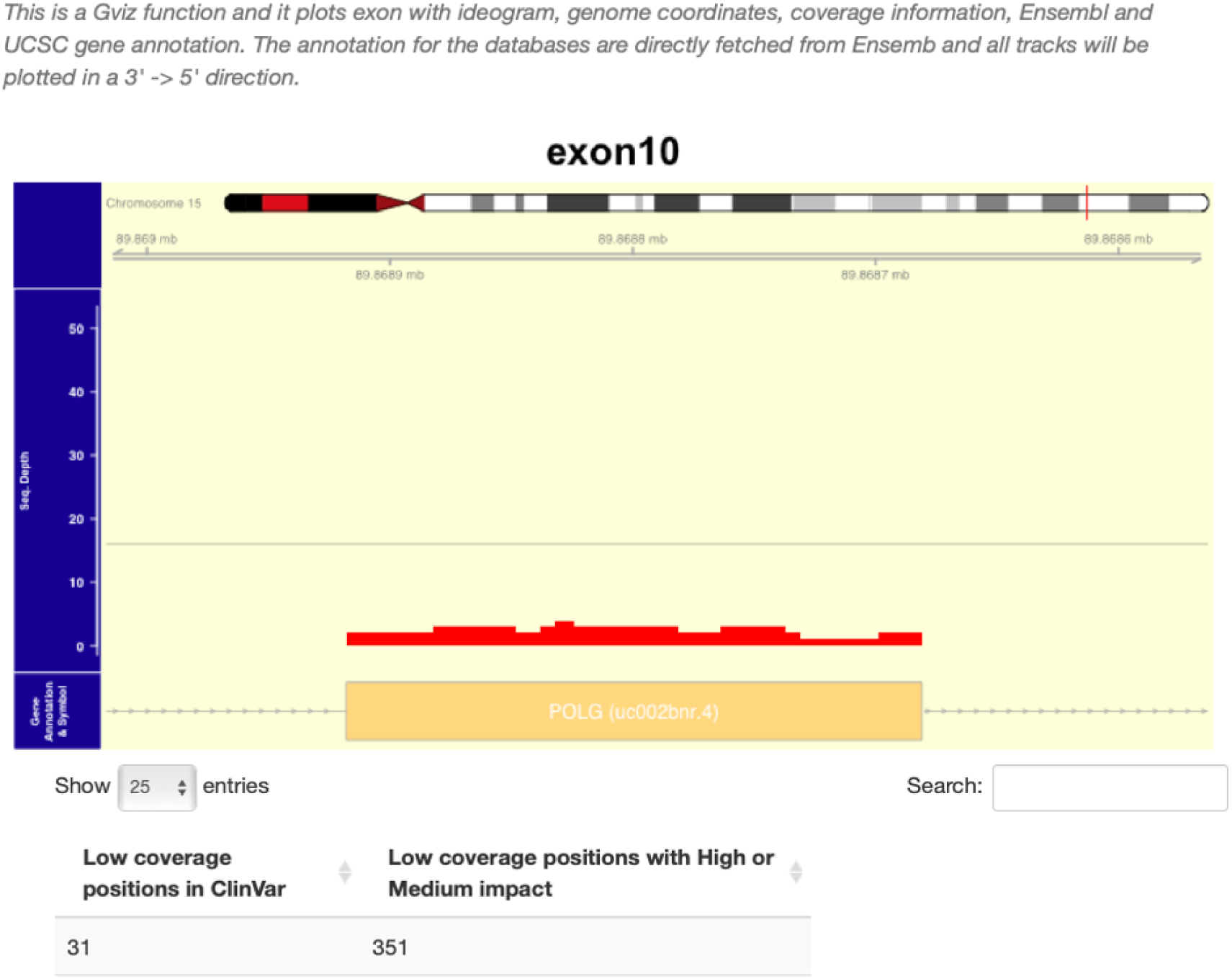
The figure shows DoC of exon 10, instead the table summarizes the number of positions known in ClinVar and with a High or medium impact.

dbNSFP-based annotation of all potential nucleotide changes across low-DoC POLG are available in `Annotation on low-coverage positions` box. The output is a downloadable table (Figure 5) displaying low-DoC positions at base-pairs level in which cells highlighted according to several criteria as high impact, clinical annotation (ClinVar), low gnomAD allelic frequency (<0.5), a damaging M-CAP score and CADD-score <20. Moreover, a low-DoC genomic position is yellow highlighted when a damaging score is found in all considered dbSNP-predictor.

**Figure 5:**
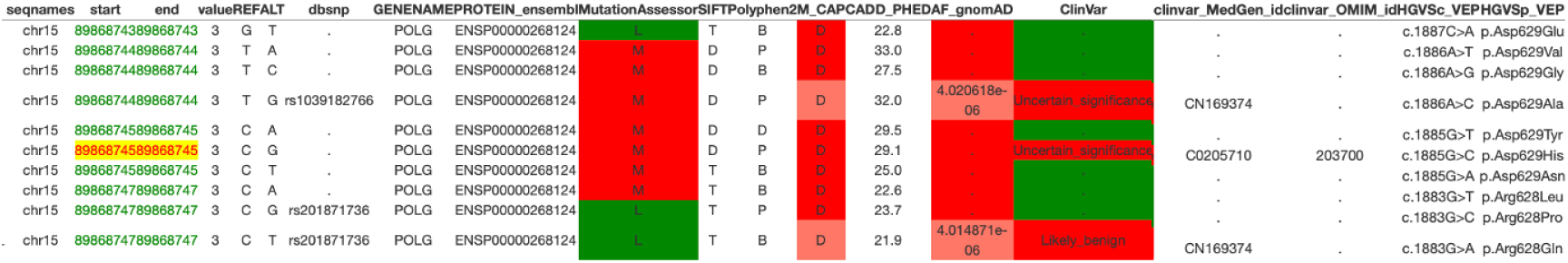
Low-DoC positions annotated with dbNSPF. dbSNP-annotation collects all consequences found in VEP-defined canonical transcripts.

However, in `Coverage Analysis` page, the default AF threshold for a variant to be annotated is 5%, then we used calculator of max AF available in `calculate AF by allele frequency app` in order to use a allele frequency based on genetic architecture of observed disease.

Analyzing functional and clinical annotation reports has allowed us to find a interesting low DoC position that hidden a probably causative variant. IGV graphical inspection had shown clinical relevant alternative allele in that low-covered position later confirmed by Sanger sequencing.

Importantly, unCOVERApp supports a binomial calculator expressing the probability that a variants is missed given its expected allelic fraction and sequencing coverage. Actually, the 20x minimum DoC threshold is reasonable for germline events where the expected fraction of variant alleles is around 0.5. Conversely, adequate DoC to detect somatic variants is paramount as that fraction can be substantially lower. unCOVERApp `Binomial distribution` page provides a simple statistical framework to evaluate if DoC is adequate to somatic variant detection. The user can set the allele fraction expected for the disease-related variant and the number of variant reads necessary to support variant calling.

In the below example, it unCOVERApp exploits binomial distribution to understand, with 95% probability, the number of reads that support a somatic variants given an `allele fraction` and the minimum number of `variant reads` required to support variant calling. The outcome in consideration box is marked in red when calculated number of reads are lower than variants reads user-defined(Figure 7), otherwise the letters are blue.

**Figure 6:**
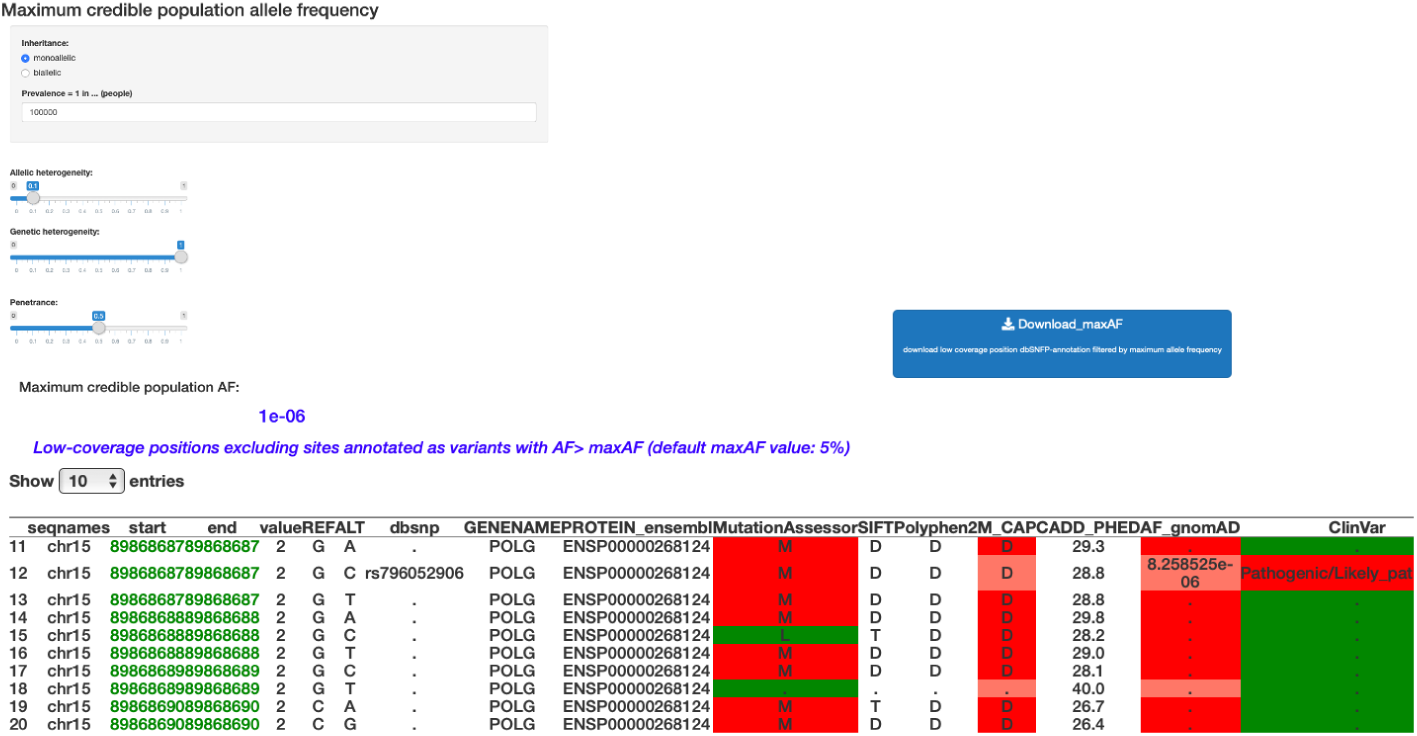
unCOVERApp allows to draw AF thresholds based on genetic architecture of condition through integration of Calculator of maximum credible population Allele Frequency.

**Figure 7:**
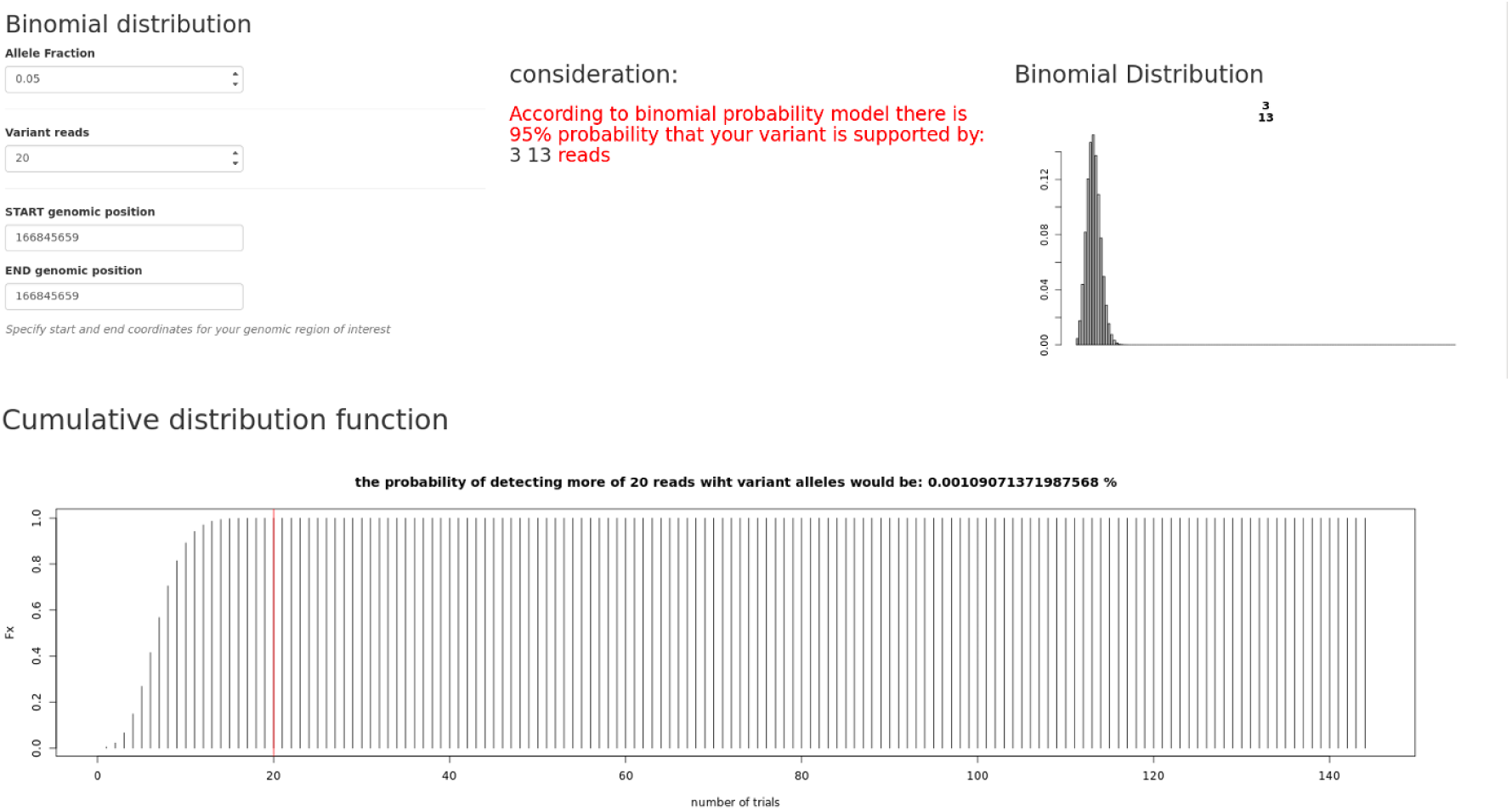
The figure shows binomial distribution analysis. The box consideration shows the number of reads that support variants based on users-defined inputs. Cumulative distribution function shows the probability of detecting less than or equal reads to the expected fraction of variant reads (probability of success) in user-defined reads (n trials).

## References

Biesecker, LG et al (2014). Diagnostic clinical genome and exome sequencing. N Engl J Med 2014;371:1170.

Casper, J, et al (2018). The UCSC Genome Browser database: 2018 update. Nucleic Acids Res. 46(D1):D762–D769.

Gargis, AS, et al (2012). Assuring the quality of next-generation sequencing in clinical laboratory practice. Nat Biotechnol. 30(11):1033–1036.

Green RC, et al (2016). Clinical Sequencing Exploratory Research Consortium: Accelerating Evidence-Based Practice of Genomic Medicine. Am J Hum Genet. 98(6):1051–1066.

Hahne, F. and Ivanek, R. (2016) Visualizing genomic data using Gviz and bioconductor. Methods Mol. Biol., 1418, 335–351.

Kumar, P et al. (2009) Predicting the effects of coding non-synonymous variants on protein function using the SIFT algorithm. Nature Protocols volume 4, pages 1073–1081 (2009)

Liu et al. (2016) dbNSFP v3.0: A One-Stop Database of Functional Predictions and Annotations for Human Non-synonymous and Splice Site SNVs. Human Mutation. 37:235–241.

Matthijs, G et al (2016). Guidelines for diagnostic next-generation sequencing. Eur J Hum Genet. 24(1):2–5.

Thorvaldsdóttir, H et al (2013). Integrative Genomics Viewer (IGV): high-performance genomics data visualization and exploration. Brief Bioinform. 14(2):178–192.

Richards S, Aziz N, Bale S, et al. Standards and guidelines for the interpretation of sequence variants: a joint consensus recommendation of the American College of Medical Genetics and Genomics and the Association for Molecular Pathology. Genet Med. 2015;17(5):405–424.

Sanghvi, RV et al (2018) Characterizing reduced coverage regions through comparison of exome and genome sequencing data across 10 centers. Genet Med. 20(8):855–866.

Whiffin, N et al. (2017) Using high-resolution variant frequencies to empower clinical genome interpretation. Genet Med. 19(10):1151–1158.

Zerbino, DR et al (2018). Ensembl 2018. Nucleic Acids Res.46(D1):D754–D761.

